# Alterations in Nutrient Availability in the Lungs During *Streptococcus pneumoniae*-Induced Pneumonia

**DOI:** 10.1101/2025.07.14.664699

**Authors:** Hansol Im, Vipin Chembilikandy, Adonis D’Mello, Madison Pearson, Hervé Tettelin, Carlos J. Orihuela

## Abstract

*Streptococcus pneumoniae* is a leading cause of pneumonia. Importantly, the extent and impact of changes in the infected airway on bacterial nutrient availability and gene expression are not known. Utilizing untargeted UPLC-ESI-MS/MS metabolomics, we comprehensively characterized the metabolic landscape in the airway across early, mid, and severe stages of pneumococcal pneumonia. This revealed that dynamic shifts in metabolites occurred during pneumonia, with an initial influx of metabolites at the early stage, followed by declines as the disease progressed. Specific host metabolic perturbations were indicative of purine dysregulation, cellular stress, and outright tissue injury. Levels of glucose, a known modulator of pneumococcal capsule production, were highest at early disease stage, and then declined as the disease progressed, overlaying general metabolite trends. Concurrent bacterial transcriptome profiling was performed using a NanoString nCounter custom panel of 66 genes selected for their importance to metabolism, virulence, and stress response; 9% of which had disease-stage significant differences in gene expression. This analysis revealed remarkably high expression of *spxB*, the gene encoding pyruvate oxidase, at the severe stage of pneumonia compared to the mid-stage pneumonia, consistent with a drop in glucose levels and indicative of a shift towards mixed fermentation and the increased production of hydrogen peroxide. Our study improves our understanding of how pneumococcal infection alters the lung environment, driving profound metabolic shifts that, in turn, influence bacterial phenotypes. This detailed understanding of host-pathogen metabolic interactions offers valuable insights into novel therapeutic strategies.

## Introduction

*Streptococcus pneumoniae* (*Spn*) is an obligate human pathogen and the leading cause of community-acquired pneumonia. While colonization by this Gram-positive pathobiont is most often asymptomatic, *Spn* can also cause life-threatening diseases such as pneumonia, bacteremia, sepsis, and meningitis, particularly in individuals with immature, deficient, or compromised immune systems (1–3). Notably, *Spn* exhibits anatomical site-specific gene expression profiles (4, 5). For instance, the transcriptome of *Spn* in the nasopharynx displays a distinct profile compared to that of bacteria found in the bloodstream, and pneumococci in the bloodstream have a different transcriptome profile than those found within invaded organs, such as the heart (5). Along such lines, prior reports by our group have demonstrated that carbohydrate availability present in specific anatomical sites is a key factor influencing pneumococcal gene expression, and in turn, this directly influences site-specific fitness within the host (4, 6). For example, under glucose restricted conditions such as in the nasopharynx, *Spn* produced less capsular polysaccharide and showed a more adhesive phenotype. Similarly, the absence of glucose in the nasopharynx has been shown to promote biofilm formation (6). In contrast, in glucose rich conditions, as encountered in the bloodstream, *Spn* grows planktonically and produces a thicker capsule which confers greater resistance against opsonophagocytosis, leading to better survival within the blood system (4, 7).

Not surprisingly, mutation of genes involved in *Spn*’s central carbon metabolism leads to impaired physiology and attenuation *in vivo* (8–10). These mutations disrupt ATP production, alter redox balance, and impact the production of essential precursors for *Spn* biology and virulence. Studies on knockouts of pyruvate oxidase (*spxB*), which convert pyruvate to acetyl phosphate, and lactate dehydrogenase (*ldh*), which converts pyruvate to lactate and back, both alter redox balance, and their deletion has been shown to result in a reduction in growth rate.

Furthermore, *spxB* and *ldh* knockouts showed significantly decreased virulence compared to the parental strain in both intranasal and intravenous infection models. Similar observations linking altered metabolism and reduced virulence have been reported for mutations in other fermentation genes such as *pfl*, *adh*, and *lctO* (11–13). Our previous report also found that knockout of fermentation genes led to altered energy production and virulence (8).

The lung environment undergoes significant changes during bacterial pneumonia due to inflammation and damage caused by bacterial factors and/or host immune responses (14–16). The response to pathogens leads to both localized and systemic inflammation, influencing tissue integrity and edema, and can, on its own, contribute to cellular damage (15). Bacterial moieties, such as pneumolysin and surface molecules, directly contribute to alveolar damage in the lung, leading to the release of nutrients from dying host cells, vascular and serum leakage, immune cell infiltration, and fluid influx in the air sacs (17, 18). A recent study highlighted that pneumococcal infection significantly alters metabolite composition compared to naïve controls, specifically identifying significant changes in branched-chain amino acid (BCAA) levels (19). Not surprisingly, *Spn* has evolved to exploit localized cellular damage, bind to dying host cells, and co-opt normally intracellular host cell factors for its advantage (20, 21). The overall findings are consistent with the observation that pneumonia caused by most virulent strains of *Spn* is characterized by significant reconstruction and inflammation of the pulmonary system (22, 23).

Lung injury during pneumonia and the inflammation needed to contain and eradicate the pathogen are continuous processes that develop over time (24, 25). Along with this, changes in nutrient availability occur within the airway. Therefore, it is reasonable to propose that the physiology and virulence profiles of *Spn* shift in response to changes in pulmonary environments during the course of infection. Yet, there are only a limited number of studies that explore the temporal kinetics between metabolite shifts and their consequences on *Spn* physiology and virulence during pneumonia. To address this lapse, this study investigated the temporal shift of lung metabolites during pneumococcal infection. Moreover, characterized overall bacterial transcriptome shift responses to this dynamic microenvironment. Using metabolomics and transcriptomics, we elucidate temporal changes in the airway as result of infection and in turn link them to associated virulence shifts, providing mechanistic insights into the pathogenesis of pneumonia.

## Results

### *Spn* infection and mediated inflammation alter the lung environment with a drastic metabolite shift

The metabolite shifts during pneumococcal pneumonia were evaluated by examining distinct stages of disease progression in mice following intratracheal challenges. As outlined in the schematic (Figure 1A), we categorized pneumonia into three stages based on previously validated health scoring criteria (26). Bronchoalveolar lavage fluid (BALF) was collected from mice at each stage of pneumonia, and detailed metabolite composition was measured. A 3D-principal component analysis (PCA) revealed that metabolite profiles during pneumonia were distinct from those of uninfected controls (Figure 1B). Notably, there was a clear separation between the early and severe stages of pneumonia, whereas the BALF recovered from mid-stage pneumonia appeared to intersect with both stages, suggesting a transitional profile. Quantitative analysis of total metabolites supported the 3D-PCA findings, showing meaningful divergence from the uninfected control group (Figure 1C). The early stage of pneumonia exhibited the highest level of metabolite infiltration, which gradually declined as the disease progressed. This stage-dependent trend became more apparent in the heatmap analysis (Figure 1D), which illustrated a progressive shift in metabolite profiles across the different stages of pneumonia relative to the uninfected group. Subsequently, we performed a Variable Importance in Projection (VIP) analysis to identify the top 10 metabolites contributing to group separation in both negatively and positively charged ion profiles. In both datasets, we observed strong accumulation of metabolites during the early stage of pneumonia (Figures 1E and 1F). In the negative ionized metabolite VIP profile, xanthine, adenosine diphosphate (ADP), and guanosine monophosphate (GMP) showed VIP scores exceeding 4.5, indicating strong separation among the infection stages. Inosine monophosphate (IMP) ranked fourth, followed by metabolites such as succinate, tyrosine, phenylalanine, uridine diphosphate-N-acetyl-galactosamine (UDP-GalNAc), and uridine monophosphate (UMP), all of which had VIP scores ranging from 2.5 to 3.0 (Figure 1E). In the positive ionized metabolite VIP profile, O-acetylcarnitine and adenosine-monophosphate (AMP) exhibited VIP scores above 5.5, making them the strongest contributors to group separation. Additional metabolites such as nicotinamide, L-carnitine, dihydrouracil, and hypoxanthine had VIP scores above 4.0. Adenosine, tyrosine, GMP, and nicotinate also showed significance in the VIP results, with VIP scores ranging from 2.5 to 3.0 (Figure 1F). Notably, several of these top-ranked metabolites—including xanthine, succinate, and multiple amino acids—are known markers of inflammation and tissue breakdown. Their elevated abundance suggests pneumonia induces dynamic metabolite infiltration associated with tissue damage and immune responses. Immunofluorescent imaging further supported the notion that progressive changes in lung inflammation and tissue damage occur (Figure 2). As pneumonia advanced, bacterial burden consistently increased, confirmed by both CFU quantification and measurement of immunofluorescence in lung sections from mice following the detection of bacteria with labeled capsule antibody (Figure 2A and 2B). Host tissue damage, specifically necroptosis, a form of programmed cell death attributable to the bacterium’s pore-forming toxin pneumolysin, was quantified using the marker MLKL as previously described (27), and also accumulated with disease progression (Figure 2C). All raw results of metabolome analysis are available in Supplementary Table 1 and 2, which contain metabolome results of negatively charged and positively charged, respectively.

**Figure 1.**
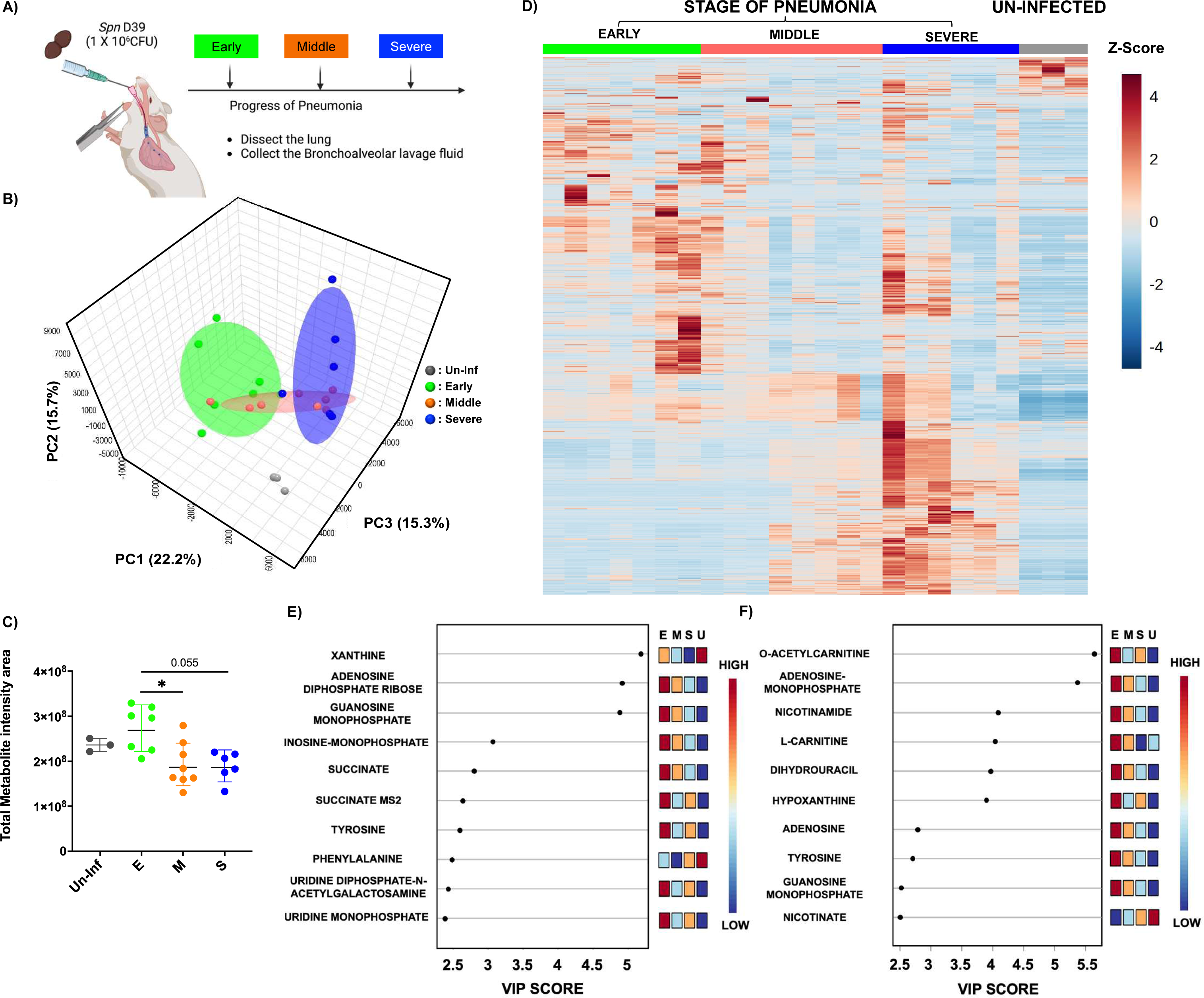
UPLC-ESI-LC-MS/MS Metabolite Analysis of Bronchoalveolar Lavage Fluid from Mock and *Spn*-Infected Mice. A) Schematics of experimental design for the study. Bronchoalveolar lavage fluid was extracted at different stages of pneumonia. B) 3D-PCA plot of metabolite compositions. C) Total metabolite intensity in bronchoalveolar lavage fluid from different stages of pneumonia. (E: Early stage of pneumonia, M: Middle stage of pneumonia, S: Severe stage of pneumonia, and Un- inf: Un-infected control). D) Heat map analysis of entire metabolites from different stages of pneumonia. E) VIP (Variable Importance in Projection) chart of top 10 metabolites from Negatively ionized mode analysis. F) VIP chart of top 10 metabolites from Positively ionized mode analysis. Significance was determined by one-way ANOVA. *, p<0.05.

**Figure 2.**
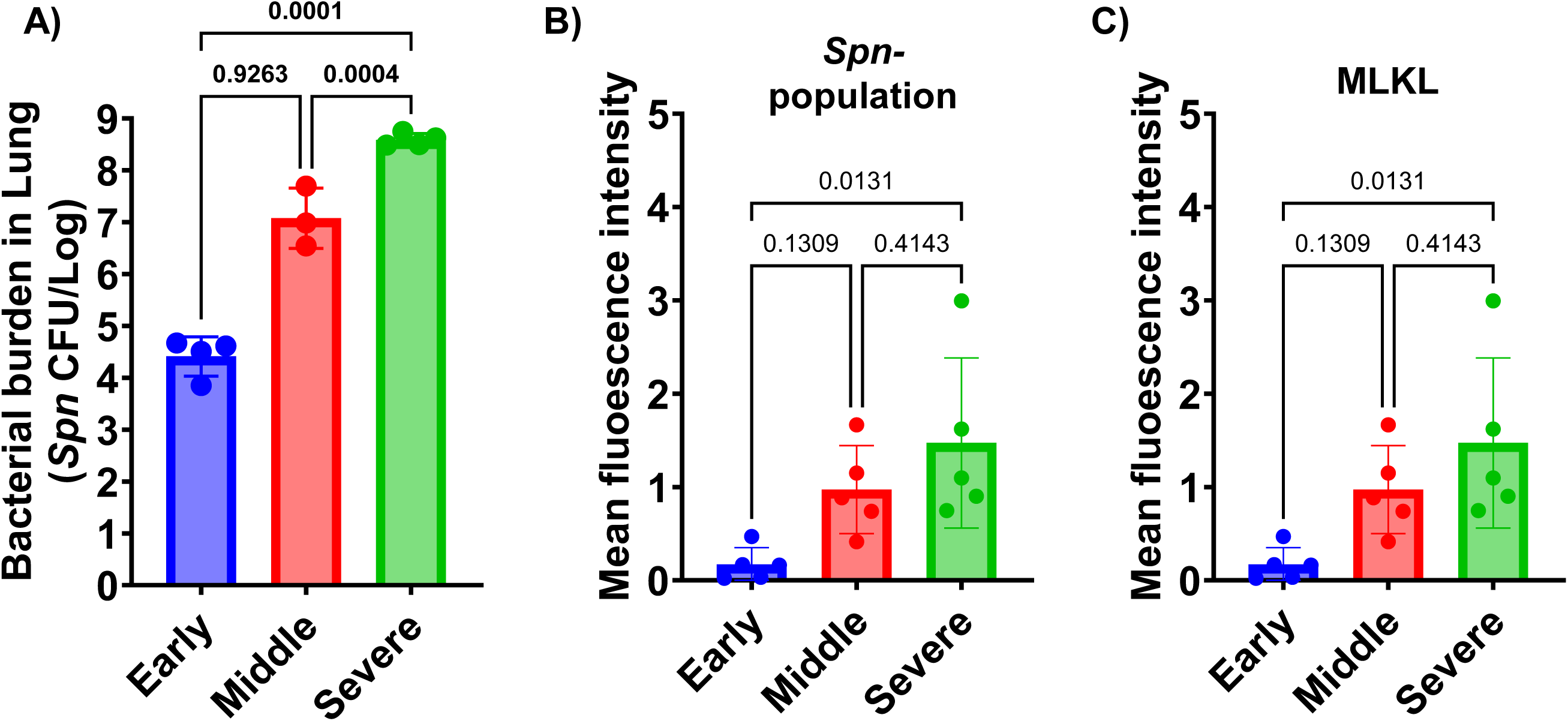
Bacterial Burden (CFU in Lung) and Microscopic Analysis Results. A) Bacterial burden in the lungs at different stages of pneumonia. B) Mean fluorescence intensity analysis of capsule-stained *Spn* within the lung section. C) Mean fluorescence intensity analysis of MLKL (mixed lineage kinase domain-like), a marker of necroptosis, across lung tissue sections from different stages of pneumonia. Significance was determined by one-way ANOVA, with p-values shown directly on the graph.

### As pneumonia progresses, metabolite profiling reveals unique metabolome profiles

To further investigate these changes, we performed Mummichog pathway enrichment analysis, which identifies biologically relevant pathways associated with altered metabolites compared to the uninfected control (28). Positively and negatively ionized metabolites were analyzed separately, revealing distinct pathway enrichment patterns across different stages of pneumonia. In the negative ion mode, we consistently observed enrichment in pyrimidine metabolism across all stages of pneumonia (Figure 3). In the early stage, 10 metabolites were significantly associated with the pyrimidine metabolism pathway (p < 0.05), and 6 with the glutathione metabolism pathway (p = 0.051). During the mid-stage of pneumonia, 65 metabolites were enriched across 11 pathways. Notable pathways included phenylalanine metabolism (6 hits), pyrimidine metabolism (12 hits), arachidonic acid metabolism (22 hits), phenylalanine/tyrosine/tryptophan biosynthesis (3 hits), and riboflavin metabolism (3 hits). In the severe stage, arachidonic acid metabolism showed the most prominent enrichment, with 22 associated metabolites approaching statistical significance (p = 0.06), suggesting a strong trend, in addition to continued enrichment in pyrimidine-related metabolites. In the positive ion mode analysis, dynamic and extensive enrichment profiles were also observed across the different stages of pneumonia. In the early stage of pneumonia, no pathways reached statistical significance. However, we observed trending results in D-glutamine/glutamate metabolism, pyruvate metabolism, and glutathione metabolism, with 5, 3, and 4 significant hits, respectively (p = 0.061, 0.067 and 0.069). In mid-stage pneumonia, 26 metabolites were significantly enriched across 3 pathways. The most distinct pathways included tyrosine metabolism (12 hits), arginine biosynthesis (8 hits), and nicotinate and nicotinamide metabolism (6 hits). At the severe stage, a notable shift was observed, with 158 metabolites associated with 39 distinct metabolic pathways. Especially, key pathways included fructose/mannose metabolism (9 hits), inositol phosphate metabolism (8 hits), phosphatidylinositol signaling (6 hits), arginine/proline metabolism (9 hits), and nicotinate/nicotinamide metabolism (4 hits). Compared to the early and mid-stage of pneumonia, the severe stage displayed unique metabolite composition, indicating a remarkable reconstruction of the lung microenvironment and immune-metabolic responses at the terminal phase of pneumonia. Overall, the data revealed that pneumonia induces a significant reconstruction of the lung microenvironment, characterized by stage-specific and pathway-specific metabolic shifts.

**Figure 3.**
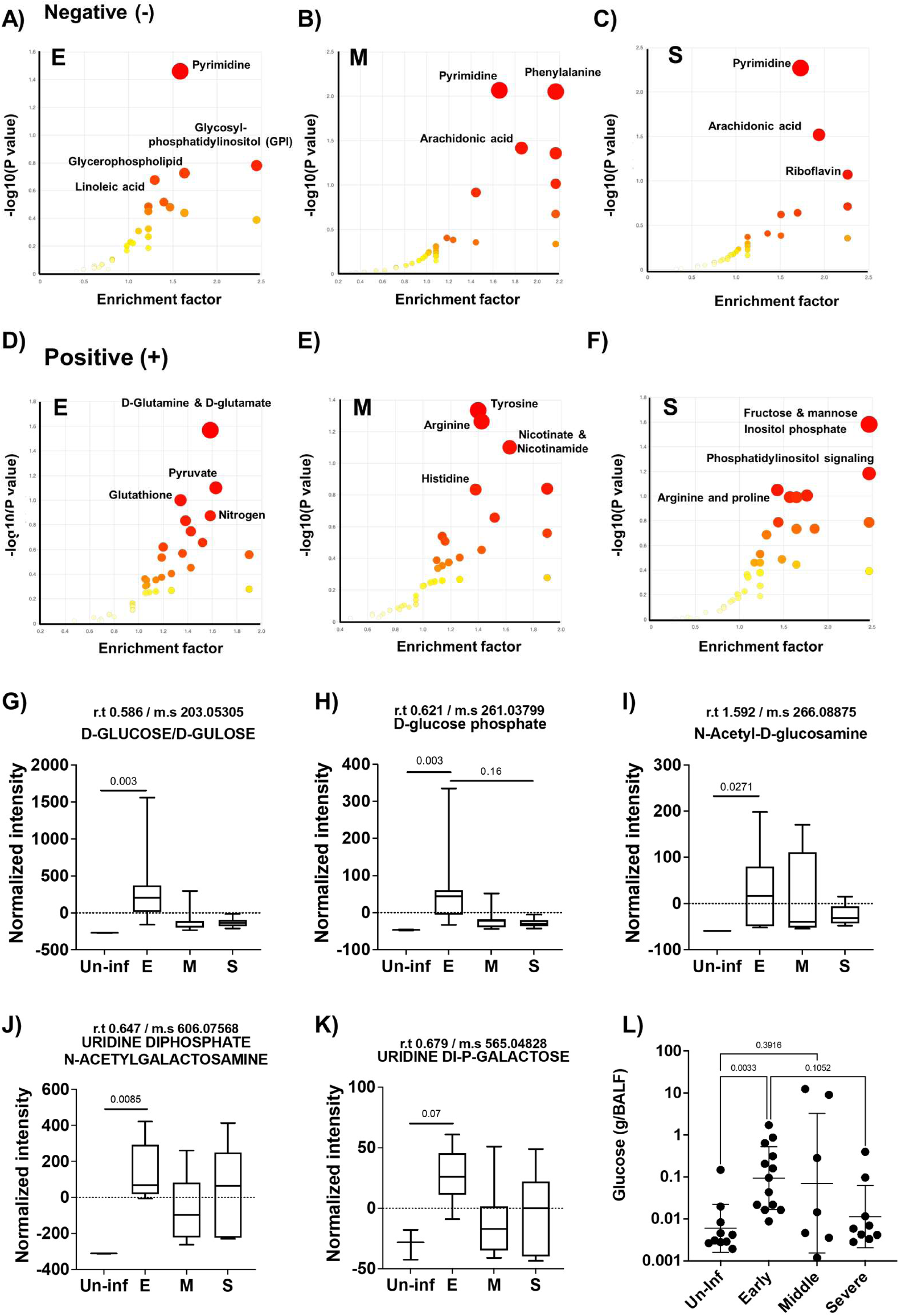
Mummichog pathway and network analysis and carbohydrate metabolite analysis of bronchoalveolar lavage fluid (BALF) from different stages of pneumonia. A-C) Negative mode mummichog analysis and metabolite enrichment results, and D-F) Positive mode mummichog analysis and metabolite enrichment results across different stages of pneumonia. Mummichog analysis was performed using Metaboanalyst 5.0 as described in methods. For the analysis, different stages of pneumonia were compared to bronchoalveolar lavage fluid from uninfected mice. (E: Early stage of pneumonia, M: Middle stage of pneumonia, S: Severe stage of pneumonia; n=7 for E, n=8 for M, and n=6 for S). G-K) Results of the carbohydrate metabolite analysis. G) D-glucose/gulose levels, H) D-glucose phosphate levels, I) N-acetyl-D-glucosamine levels, J) Uridine di-phosphate N-acetyl-galactosamine levels, and K) Uridine Di-phosphate-galactose metabolite levels from different stages of pneumonia and uninfected control group. All samples were normalized using total recovered volume of bronchoalveolar lavage fluid. L) Glucose concentration in recovered bronchoalveolar lavage fluid from different stages of pneumonia and uninfected control group. Significance was determined by one-way ANOVA, with p- values shown directly on the graph.

### Increased levels of glucose and other carbon metabolites were observed in the early stage of pneumonia

As carbon availability is a key determinant of *Spn* physiology, virulence, and survival within the host, we specifically assessed the dynamics of carbohydrate-related carbon sources during pneumonia progression (Figure 3G–2L). Metabolomic profiling of BALF samples revealed an early spike in overall carbohydrate levels compared to uninfected controls (Figure 3G–2K). This difference was only detected during the early stage. In Figures 3G and 3H, we analyzed D-glucose and D-glucose phosphate, both of which showed significant increases in the early stage, followed by a decline as the disease progressed. In Figure 3I, N-acetyl-D-glucosamine levels significantly increased early in infection and showed a decreasing trend over time, though mid-stage pneumonia displayed some variability, suggesting a metabolic transition phase. Figures 3J and 3K depict uridine diphosphate N-acetyl-galactosamine and uridine diphosphate galactose, respectively.

These metabolites also showed significantly elevated or trending increases during the early stage. Although the differences were not statistically significant, the average metabolite levels in BALF from the severe stage of pneumonia were higher than those observed in the mid-stage of pneumonia BALF. To validate these findings, we quantified glucose levels in extracted BALF samples. Consistent with our metabolomic data, glucose concentrations were approximately 14-fold higher in the early stage of pneumonia compared to uninfected controls and then declined as pneumonia progressed from the early to the severe stages. Although glucose levels in mid- and severe-stage samples remained slightly elevated relative to controls, no statistically significant differences were observed. Thus, available carbohydrates rapidly escalate during early infection and then decrease but remain elevated compared to uninfected as the infection proceeds.

### Shifted metabolite profiles of lung during pneumonia alter the *Spn* transcriptome

Transcriptome analysis using a custom-made Nanostring nCounter panel of probes interrogated the expression levels of 66 target genes related to bacterial metabolism, stress responses, virulence factors, and surface proteins (Supplemental Table 3). *Spn* CFUs in mouse lungs from early stages of pneumonia were insufficient for robust detection of transcripts, therefore differential gene expression analyses were performed for the severe stage compared to the mid-stage. Among the 66 genes, six genes were differentially expressed between the two different stages of pneumonia. At the severe stage of pneumonia, we found six genes to be highly expressed compared to the mid-stage, which included SP_0730 (*spxB*), and SP_0366 (*aliA*). In the mid-stage of pneumonia, we observed high expression of SP_2106 (*malP*), SP_0648 (*bgaA*), SP_0966 (*pavA*), and SP_1675 (a kinase group gene) as presented in Figure 4 and Supplementary table 3. Among these six genes, we observed relatively higher overall gene expression of *spxB* and *aliA*, with significant differences between the two different stages. Intriguingly, we observed significantly higher standard deviation in the mid-stage of pneumonia which was similarly observed in metabolome analysis.

**Figure 4.**
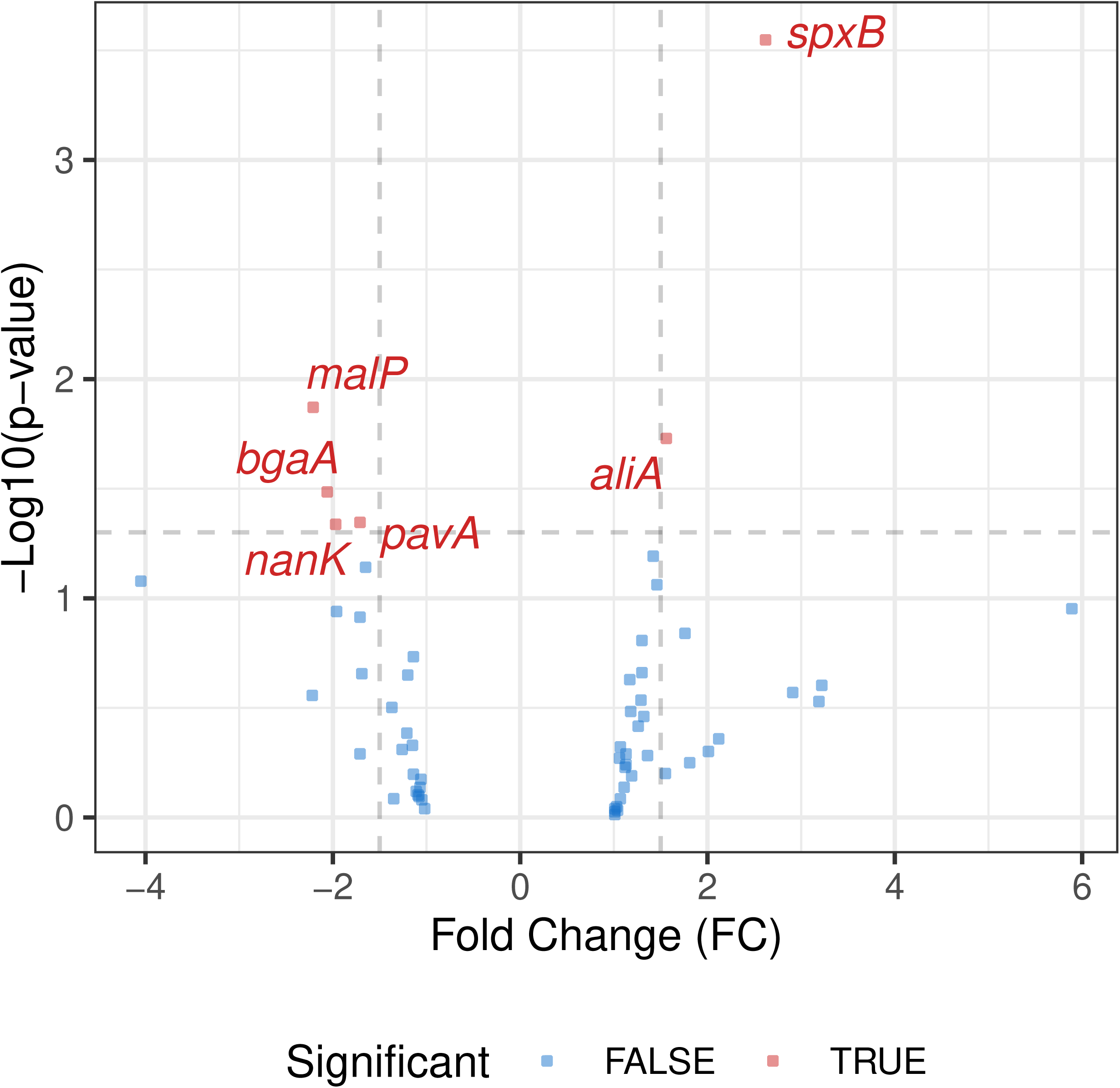
Volcano plot of Nanostring nCounter gene expression analysis between mid-stage and severe stage of pneumonia. Analysis was performed using remaining genes that met significance cutoffs (p-value ≤0.05 & absolute fold change ≥1.5). Genes below the average counts values (<50) in either condition were not included for the analysis.

## Materials and Methods

### Bacterial strains

The strain used in this study is *S. pneumoniae* D39 (5). Bacteria was grown in Todd-Hewitt broth with 0.5% yeast extract (THY), or on blood agar plates (Remel), in a humidified atmosphere at 37°C with 5% CO2. Note that for all analyses, we used bacteria growing in early exponential phase, corresponding to an optical density at 620 nm (OD620) of 0.25 to 0.40.

### Ethics statement

All mouse experiments were reviewed and approved by the Institutional Animal Care and Use Committee at The University of Alabama at Birmingham, UAB (protocol no. IACUC-21851). Animal care and experimental protocols adhered to Public Law 89–544 (Animal Welfare Act) and its amendments, Public Health Services guidelines, and the Guide for the Care and Use of Laboratory Animals (29).

### In vivo infection model

For the pneumonia model, 1 × 10^6^ CFU of *Spn* D39 in 100 µL was inoculated intratracheally as described previously (4). Different stages of pneumonia (Early, Middle, and Severe) are determined based on health index score as described in previous report (26). Bacterial burden in serum was measured using tail-snip blood collection at different stages of health index. At each stage of pneumonia, mice were sacrificed, and harvested lungs for analyzed for imaging and transcriptome analysis. Lung samples were processed for both paraffin and frozen section by different dissection. Bronchoalveolar lavage fluid collection was done separately to avoid physical stress on lung samples. 3 ml of PBS was loaded to airway system and recovered. Bacterial burden was measured using BALF and left BALF was stored at –20°C for further analysis.

### Metabolome analysis of Bronchoalveolar lavage fluid

Extracted bronchoalveolar lavage fluid samples are mixed with chloroform/methanol in a 1:2 ratio (v/v). 500 µl of samples were mixed with 1.9 ml of chloroform/methanol (1:2 ratio v/v) and vortexed. Afterward, 625 µl of chloroform and 625 µl of ddH2O were added sequentially with vortexing. Mixed samples were centrifuged at 3,000 rpm for 10 minutes, and we collected the top aqueous/methanol layer and placed it in a new test tube. The aqueous phase was dried under nitrogen gas. Dried supernatants were re- suspended in 100 µl of 0.1 formic acid in ddH2O and injected 20 µl of sample volume on the SCIEX 5600 Triple-Tof mass spectrometer for the analysis. UPLC-ESI-MS/MS data-dependent analyses in both positive and negative mode ionization. For data analysis, we used Metaboanalyst 5.0 for metabolome results in both pathway analysis and statistics. Statistics for metabolome analysis with Metaboanalyst were conducted as described previously(28). Pathway analysis was also conducted using same software.

**Immunofluorescence assay.** Tissue-Tek O.C.T. Compound (Sakura Finetek USA) embedded section was processed to remove the resins using cold acetone (10 min), cold 70% ethanol (5 min) and washed with PBS (Fisher scientific, USA) for 3 minutes. After OCT compound was removed, each slide was permeabilized using PBS-triton X (0.1%) for 5 minutes. Permeabilized sections were rehydrated with PBS for 5 min and blocked with 5% BSA in PBS for 30 minutes. Primary antibody, mouse anti-Spn serotype 2 (1:1000 dilution) or rabbit anti-MLKL (1:1000 dilution), was added to the slides and incubated overnight at 4 °C. For secondary antibody, we used Cy3 goat anti-mouse IgG (Invitrogen-USA) and Alexa Fluor 594 donkey anti-rabbit IgG conjugate for visualization. Secondary antibody was added 1:1000 diluted concentration to ensure the high signal, and incubation was accomplished for 3 hours at room temperature. The slides were washed with PBS (3 min) after incubation. The slides were counterstained with DAPI (NucBlue-Invitrogen, USA) at a concentration of 2 drops in 1 mL PBS. The sections were mounted with Fluoromount™ Aqueous Mounting Medium (Sigma, USA), and covered with coverslips for microscopy. Images of lung sections were captured using a Leica LMD6 microscope equipped with a DFC3000G monochrome camera. Image stitching of whole immunofluorescence microscopy-stained lung sections was performed using Leica LASX software. The fluorescence intensity was measured using FIJI/Image J.

### RNA isolation

Mouse tissue samples in RNAprotect were centrifuged at 5,000 g for 5 minutes and the supernatant was discarded. Pellets were then resuspended with 1 mL of cold PBS and spun again at 5,000g for 5 minutes to wash the pellet. Supernatant was discarded. Pellets were then resuspended in 100 μL of lysis buffer (10 μL of mutanolysin, 20 μL of proteinase K, 30 μL of lysozyme, 40 μL of TE buffer) followed by mechanical disruption using a motorized pestle for 30 seconds. Samples were then incubated for 10min at room temperature with vortexing every 1-2mins. After incubation, 600uL of Qiagen buffer RLT was added and samples disrupted again using a motorized mortar and pestle for 30 seconds. The lysate was then homogenized using a Qiagen Qiashredder column spun at 16,000 g for 2 minutes. RNA was then precipitated from the resultant flowthrough and captured on the RNeasy Mini Kit columns with DNase treatment on column (Qiagen protocol). Extracted RNA was quantitated using a Bioanalyzer.

### Transcriptome profiling using NanoString

NanoString differential gene expression analysis was performed in nSolver comparing severe to mid-stage samples. Genes having average counts values (>50) in either condition were deemed above noise. Volcano plots were created with remaining genes that met significance cutoffs (p-value <=0.05 & absolute fold change >=1.5).

## Discussion

The host microenvironment surrounding bacterial pathogens undergoes significant alterations during infection, profoundly influencing host and bacterial physiology, as evidenced by numerous studies (19, 30). The nature of the host environment is dynamic and influenced by factors such as bacterial-mediated damage, inflammation, aging, and genetics (31, 32). Infections and their associated damage are major drivers of acute/rapid host structural changes in the airway, inducing substantial remodeling of inflamed tissues and systemic responses (33, 34). Similar to the report by Sender et al. on co-infection models with influenza, which observed high concentrations of metabolites that infiltrated from vascular leakage and an increased bacterial burden, we also found rapid and significant changes in metabolite profiles following pneumococcal infection (30).

One of our most interesting findings was the highly differentiated metabolite compositions observed throughout the course of pneumonia. While previous studies noted higher glucose concentrations in lung inflammation, particularly in COPD patient sputum and influenza-infected lung environments (35, 36), our study revealed a significant influx of metabolites, beyond that of carbohydrates, predominantly at the early stage of pneumonia, which then decreased as pneumonia progressed. The development of pneumonia in our model coincided with severe lung damage, indicating intense inflammation, as our previous report previously demonstrated (27). This distinct metabolome pattern, characterized by initial high enrichment followed by a decline, contrasted with findings in certain other inflammation models.

The distinct metabolome profiles of hypoxanthine and xanthine in the lung metabolome highlighted dynamic alterations in host purine metabolism during pneumococcal pneumonia. We observed a marked accumulation of hypoxanthine during the early stage of pneumonia, followed by a decline in later pneumonia stages. This initial surge in hypoxanthine suggests early tissue damage and ATP depletion, which are known consequences of purine catabolism under hypoxia or vigorous innate host immune responses (37). While xanthine oxidase (XO) activity is known to increase significantly under inflammatory and hypoxic conditions, leading to reactive oxygen species (ROS) generation (38), we observed an intriguing pattern for xanthine. Xanthine was highly abundant in uninfected controls but drastically decreased during pneumonia progression. We initially expected an increase in xanthine given the heightened XO activity. However, our findings, supported by the concurrent accumulation of guanosine monophosphate (GMP), adenosine monophosphate (AMP) and adenosine-di-phosphate ribose (ADP-ribose), offer a compelling explanation. The accumulation of GMP, AMP and ADP-ribose reflects a state of significant cellular stress and purine dysregulation (39). The elevation of these purine nucleotides indicates dysfunction of purine metabolism, while ADP-ribose accumulation specifically points to increased NAD+ consumption, likely due to activation of stress-response pathways such as PARPs (Poly [ADP-ribose] polymerases), which are known to be activated during inflammation and DNA damage (40). These findings, coupled with high hypoxanthine levels and decreased xanthine concentrations, support a widespread perturbation of host purine metabolism in the early phase of pneumonia, providing abundant substrates that are then rapidly processed by the inflammation-induced XO, likely driving the rapid conversion of xanthine to uric acid and contributing to the overall inflammatory milieu (38).

Similarly, we observed strong accumulation of inosine monophosphate (IMP), succinate, tyrosine, O-acetylcarnitine, nicotinamide, L-carnitine, and dihydrouracil. IMP, a precursor for AMP and GMP, regulates intracellular purine metabolism, and its accumulation alongside other purine metabolites further indicates host nucleotide dysregulation (41). The accumulation of succinate and altered tyrosine levels also support acute tissue injury and metabolic alterations associated with pneumonia, potentially reflecting a bottleneck in the TCA cycle and broader damage to metabolic systems within the organ (42, 43). Furthermore, increases in pyrimidine metabolites, including uridine diphosphate-N-acetyl-galactosamine (UDP-GalNAc), uridine monophosphate (UMP), and dihydrouracil, corroborate extensive nucleic acid turnover and cellular damage (44). Complementing these specific metabolite observations, our Mummichog pathway enrichment analysis also showed broader stage-specific and pathway-specific metabolic shifts by integrating findings from both positive and negative ion modes. We observed strong and consistent enrichment of pyrimidine metabolism pathways, correlating with our findings on specific uridine metabolites. Additionally, antioxidant-related metabolic pathways, including D-glutamine/glutamate metabolism and glutathione metabolism, primarily in the positive ion mode, were also notably enriched at the early stage of pneumonia (45).

The most intriguing observation across the majority of our metabolites was the pattern showing increases at the early stage and decreases as pneumonia developed. We propose multi-factorial reasons for this dynamic. First, given that *Spn* can consume diverse nutrients and possesses associated metabolic pathways, an increased bacterial burden at later stages could relate to the decreased metabolite concentrations as the pathogen scavenges available host-derived moieties (46, 47). Second, vigorous consumption and remodeling by host immune cells are also likely contributors. Research indicates that immune cells undergo significant metabolic shifts during acute infection, demanding substantial nutrient sources for activation and function (48). Thus, a vigorous host immune response could be another factor associated with this unique biphasic metabolite pattern. However, drawing definitive conclusions necessitates further mechanistic studies to precisely disentangle the contributions of host and pathogen metabolism.

Reflecting the dynamically altered metabolite landscape, we observed a notable shift in the bacterial transcriptome. A total of 9% of the genes on our Nanostring panel reached significant differential expression and this difference was most likely representative of the transition state between early and severe infection profiles. An interesting finding was the high expression of *spxB* (SP_0730) at the severe stage of pneumonia, as pyruvate oxidase is known to be highly expressed under nutrient-limited conditions and be an indicator of mixed fermentation (4). A byproduct of SpxB-mediated fermentation is hydrogen peroxide, which in turn has been shown to be cytotoxic to lung cells. Moreover, elevated *spxB* expression has been shown in our hands to coincide with lower levels of bacterial capsule production and enhanced bacterial adherence. Notably, recent work by Alibayov et al. reported that hydrogen peroxide from *Spn* oxidizes the hemoglobin present in the alveoli, releasing hemes (49). Given our observation of a higher bacterial burden in the lung, these transcriptional changes support the hypothesis that the decreased availability of host nutrients at the severe stage creates a challenging, nutrient-limited environment for *Spn –* but also an environment that *Spn* can adapt to at the host’s detriment. Interestingly, for *bgaA* (SP_0648), encoding a putative beta-galactosidase important for host glycan utilization (50), we observed an opposite trend with a difference of only around 2-fold, a magnitude lower than anticipated based on our previous report that compared CDM-B vs CDM-N (4). This modest change possibly reflects the diversified conditions, and bacterial metabolic adaptations present across the mid-stage of pneumonia.

Critically, our observations herein that *Spn* is modulating its gene expression in response to the host condition, particularly that associated with damaged or dying host cells, are consistent with our prior findings which showed *Spn* specifically co-opts dying host cells to its advantage (20). Along such lines, *Spn* binds to dying host cells via PspA, which binds to surface-associated GAPDH on dying host epithelial cells and co-opts released lactate dehydrogenase from dying cells to promote its virulence. These traits, along with the metabolic adaptations described herein, have likely evolved such that the pneumococcus can take advantage of inflammatory conditions in the nasopharynx, such as following viral co-infection, and promote its transmission (51). That they also function to cause pneumonia is likely incidental.

In conclusion, our findings demonstrate that pneumococcal infections dynamically alter the lung microenvironment, resulting in stark changes in available metabolites. These metabolic shifts, in turn, influence bacterial behavior and contribute to unique transcriptome profiles that can benefit bacterial survival, a finding consistent with our previous report (4, 5). We believe our study expands our understanding of how host metabolites and nutrient availability shifts during the course of pneumonia and its influence on bacterial behavior. This detailed understanding provides valuable targets for therapeutic intervention, as our previous report suggested such targeting of the redox system in *Spn* (8).

## Acknowledgements.

This project was funded by NIH AI114800, AI156898, and AI172796 to CJO. Funds provided during Merck Investigator Studies Program (MISP) agreement #60280 to HT supported the development and purchase of the NanoString nCounter probe panel used in this study. We also acknowledge essential support provided by members of the Institute for Genome Sciences’ Maryland Genomics core. For UPLC-ESI-MS/MS, we acknowledge great support provided by Dr. Stephen Barnes and his member to support the study.

## Author contribution

HI and CJO conceptualized the ideas and designed all experiments. HI, VC, AD, HT, and CJO conducted a review for the article. HI, VC, and MP conducted *in vitro* and *in vivo* work for the project. HI and VC conducted image analysis for the manuscript. HI, AD, HT and CJO conducted transcriptome analysis for the manuscript. HT and CJO provided resources for the project.

**Supplementary Table 1.** UPLC-ESI-LC-MS/MS analysis results of negatively charged metabolites.

**Supplementary Table 2.** UPLC-ESI-LC-MS/MS analysis results of positively charged metabolites.

**Supplementary Table 3.** NanoString nCounter results of *S. pneumoniae* (*Spn*) differential gene expression comparison between mid-stage and severe stage pneumonia.

